# Positive selection of a starch synthesis gene and phenotypic differentiation of starch accumulation in symbiotic and free-living coral symbiont dinoflagellate species

**DOI:** 10.1101/2025.03.26.645627

**Authors:** Yuu Ishii, Shunsuke Kanamori, Ryusaku Deguchi, Masakado Kawata, Shinichiro Maruyama, Takashi Yoshida, Ryoma Kamikawa

## Abstract

Symbiosis is a basis for species diversification through interactions between organisms. In tropical and subtropical oceans, dinoflagellate symbionts belonging to the family Symbiodiniaceae, including the genus *Symbiodinium*, support the flourishment of cnidarian hosts, including corals, and thereby the ecology of oligotrophic oceans through their photosynthate carbon transfers. Although the genus *Symbiodinium* includes both free-living and symbiotic species, the detailed genetic background of their lifestyle differences remains unclear. In this study, we identified candidate genes involved in the evolutionary acquisition or maintenance of symbiosis in *Symbiodinium* spp. by detecting genes that have undergone positive selection during symbiotic and free-living lifestyle diversification. Using multiple *Symbiodinium* genomes to detect positive selection, 35 genes were identified, including a gene encoding soluble starch synthase SSY1 and genes related to metabolite secretion, which may be preferred for symbiotic lifestyles. In particular, our *in silico* analyses revealed that the SSY1 gene family has undergone extensive gene duplications in an ancestral dinoflagellate, and that the mutations detected as positive selection have occurred in the intrinsically disordered region of one of the homologs. Consistent with molecular evolution, the phenotypes of intracellular starch synthesis/accumulation were distinct between the symbiotic and free-living species of *Symbiodinium* when cultured under different pH and nitrogen conditions. These results provide molecular and phenotypic insights into symbiotic *Symbiodinium*-coral relationships.

**Significance statement:** Clarifying how coral symbiont algae of the family Symbiodiniaceae are functionally diversified is crucial for understanding the molecular mechanisms of symbiotic relationships and interactions in animal-algal symbioses. However, gene-level molecular evolution that contributes to symbiotic relationships remains unclear. Our genome-wide single-gene analyses of the genus *Symbiodinium*, comprising both symbiotic and free-living species, successfully detected several dozen protein-coding genes that underwent symbiont-specific positive selection after divergence from the free-living species in the genus. Furthermore, our cultivation experiments demonstrated differences in photosynthate-related phenotypes between the symbiotic and free-living species. Our findings provide a key to illuminating the molecular evolution involved in phenotypic changes that potentially contribute to symbiotic lifestyles in *Symbiodinium*.

## Introduction

Symbiosis is a ubiquitous phenomenon wherein an organism lives with other organisms (Dimijian 2000; Gilbert et al. 2012; McFall-Ngai et al. 2013). Thus, symbiosis drives functional and ecological diversification through the mutual exchange or one-sided exploitation of metabolites and habitat provision (Gilbert et al. 2012; McFall-Ngai et al. 2013; Yellowlees et al. 2008). Coral reefs are ocean ecosystems that are highly dependent on symbiosis (Blackall et al. 2015). Photosynthesis by unicellular algal symbionts belonging to the family Symbiodiniaceae in dinoflagellates provides most of the energy input to coral reef ecosystems (Muscatine 1990), thereby supporting not only the reef’s own prosperity, but also ecosystem diversity in oligotrophic oceans (Baker 2003; Roth 2014; Simpson et al. 2011).

While species diversification and habitat expansion of stony corals in oligotrophic shallow waters emerged approximately 240 million years ago or earlier (Frankowiak et al. 2016), species diversification of the family Symbiodiniaceae was estimated to have occurred approximately 160 million years ago, suggesting that Symbiodiniaceae diversification is associated with coral symbiosis (LaJeunesse et al. 2018). Symbiodiniaceae contains 10 clades, from A to J, which correspond to genus-level taxonomic groups, and ecological, biogeographical, and genomic characterizations have been conducted (Aranda et al. 2016; González-Pech et al. 2017; González-Pech et al. 2021a; LaJeunesse et al. 2018; Lin et al. 2015; Liu et al. 2018; Shoguchi et al. 2013; Yorifuji et al. 2021). Symbiotic (but culturable without hosts in some cases) and free-living (not detected as symbionts in nature) species are included in the family, and various combinations of animal hosts and Symbiodiniaceae symbionts have been observed, forming a many-to-many symbiotic relationship (Mies et al. 2017).

In the symbiotic phase, Symbiodiniaceae symbionts conduct photosynthesis in the habitats provided by their hosts (Davy et al. 2012; Norton et al. 1992). These habitats are acidic (Armstrong et al. 2018; Barott et al. 2015), and nitrogen levels are arguably controlled by the hosts (Davy et al. 2012; Xiang et al. 2020). Symbionts transfer more than 90% of their photosynthates to their hosts (Muscatine 1990). Sugars, lipids, and amino acids are metabolites transferred by Symbiodiniaceae symbionts to their hosts (Davy et al. 2012). Gene expression analysis of symbiotic model organisms, such as sea anemones (*Exaiptasia diaphana*) and Symbiodiniaceae, suggests that sugar transporters and sugar metabolism-related genes in hosts and symbionts are involved in the maintenance of stable symbiosis (Davy et al. 2012; Gordon & Leggat 2010; Ishii et al. 2019; Maor-Landaw et al. 2020; Mashini et al. 2022; Sproles et al. 2018). This further highlights the role of sugar transfer from Symbiodiniaceae cells to the host in the evolutionary establishment of the symbiotic relationships (Davy et al. 2012; Gordon & Leggat 2010; Ishii et al. 2019; Maor-Landaw et al. 2020; Mashini et al. 2022; Sproles et al. 2018). Two mechanisms for sugar secretion from Symbiodiniaceae have been experimentally characterized: one pathway is that photosynthates synthesized in the symbiont are transferred via certain transporters in a short time (Burriesci et al. 2012; Cui et al.; Maor-Landaw et al. 2023) and the other is that sugar is gradually secreted by degrading the Symbiodiniaceae cell wall using its own cellulase (Ishii et al. 2023). Some photosynthates are not secreted extracellularly, but accumulate intracellularly in the form of starch and lipids, and the quantity of accumulated photosynthate-derived compounds varies among culture conditions (Grant et al. 2006; Weng et al. 2014).

At the genomic level, the most well-studied genus in the family is *Symbiodinium*, formally known as clade A, which is believed to have adapted to strong and variable light conditions and is distributed worldwide (LaJeunesse et al. 2018). Notably, both symbiotic and free-living species belong to the genus *Symbiodinium* (González-Pech et al. 2021a; Quigley et al. 2017; Hansen & Daugbjerg 2009). Although molecular phylogenies have shown free-living species to be paraphyletic and deep-branching in *Symbiodinium* (LaJeunesse et al. 2018), whether the last common ancestor of the genus was symbiotic or free-living remains uncertain (González-Pech et al. 2021a). Currently, genomes of seven strains from six species are available: *S. microadriaticum* (*S. mic*) CCMP2467 and 04- 503SCI.03, *S. necroappetens* (*S. nec*) CCMP2469, *S. linucheae* (*S. lin*) CCMP2456, *S. tridacnidorum* (*S. tri*) CCMP2592, *S. natans* (*S. nat*) CCMP2548, and *S. pilosum* (*S. pil*) CCMP2461, the latter two of which are free-living (Aranda et al. 2016; González-Pech et al. 2021a; González-Pech et al. 2021b) (Figure 1).

**Fig. 1.**
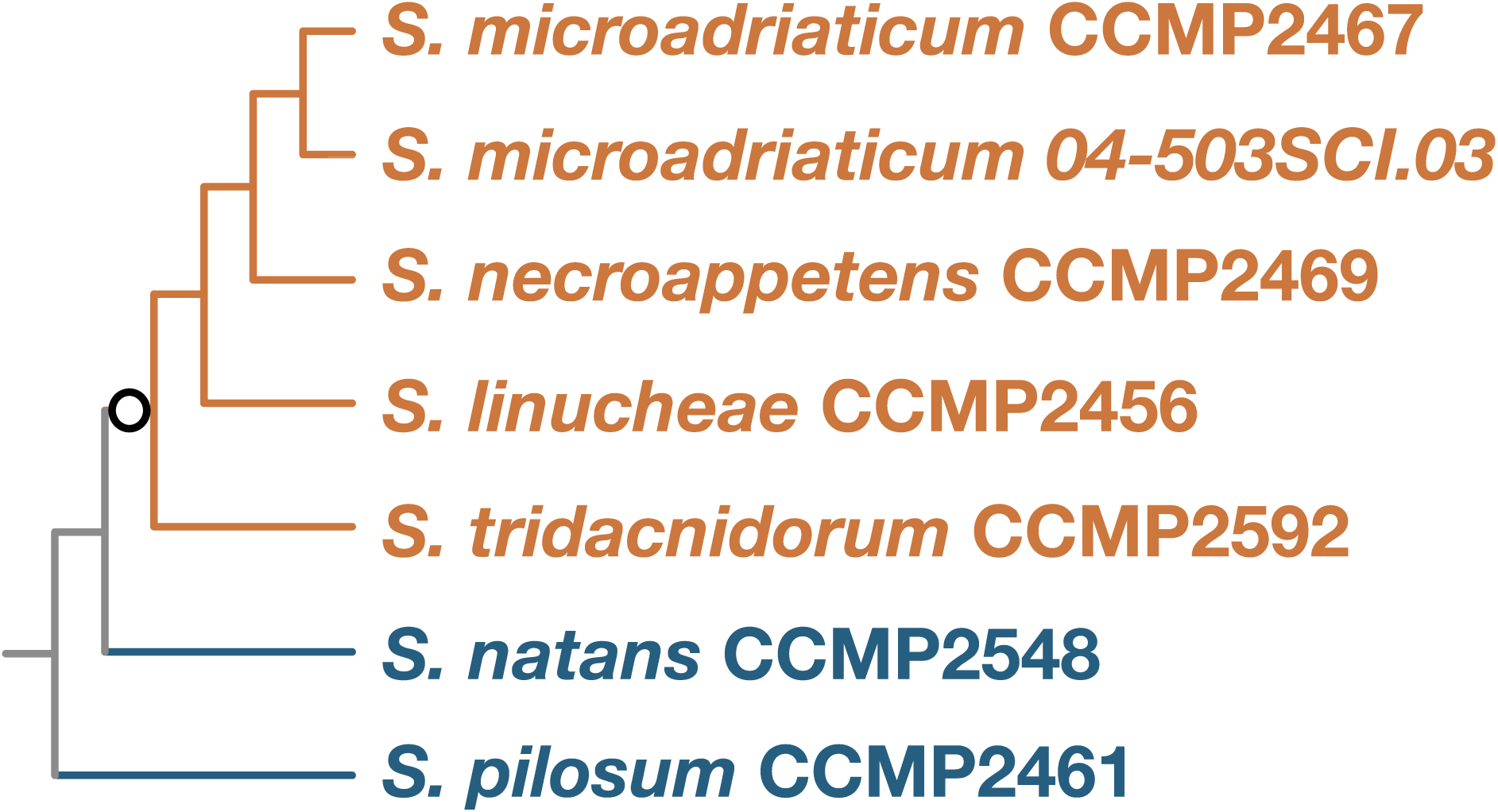
Phylogenetic relationships of seven *Symbiodinium* strains used for detecting positive selection. A cladogram of the seven *Symbiodinium* strains is constructed based on previous studies (González-Pech et al. 2021a; LaJeunesse et al. 2018). The names colored in orange and blue indicate symbiotic and free-living species, respectively. Branches descending from the edge, displayed with a circle (shown in orange), are foreground branches used to detect genes under positive selection by codeml and aBSREL.

In this study, we performed comparative analyses of all coding sequences (CDSs) in the available genomes of seven *Symbiodinium* strains from six species to detect genes that have undergone positive selection on a branch leading to the symbiotic clade after divergence from free-living clades. In addition, we experimentally evaluated phenotypic differentiation between symbiotic and free-living species of *Symbiodinium*. Overall, our results indicate that at least several dozen protein-coding genes, including those involved in starch synthesis, have been positively selected during the evolution of *Symbiodinium* and are associated with phenotypic differences in starch synthesis/accumulation between species with different lifestyles.

## Results

### Detection of positive selection during lifestyle diversification

The number of single-copy orthologs in the six species was 1,879. The phylogenetic tree topologies of the 1,309 genes were consistent with the species tree topology. These 1,309 genes were subjected to the survey of positive selection on the foreground branch leading to the exclusive common ancestor of the symbiotic species, *S. mic*, *S. nec*, *S. lin,* and *S. tri*, on the species tree (Figure 1; highlighted with open circle). The number of positively selected genes detected by codeml and aBSREL with different models was 242 and 38, respectively. Of these, 35 genes were regarded as reliable candidates for positive selection because they were detected by both codeml and aBSREL, and 18 out of 35 were not functionally annotated. Successfully annotated 17 genes (Table 1) encode soluble starch synthase 1 (*SSY1*), cholesterol transporter (*ABCA5*), peroxisomal ABC transporter (*ABCD3*), amino-acid acetyltransferase (*ARGA*), sterol 3-beta-glucosyltransferase (*ATG26*), putative sulfur deprivation response regulator (*SAC1*), calcium-dependent protein kinase 2 (*CDPK2*), sodium channel protein para (*SCNA*), tartrate-resistant acid phosphatase type 5 (*PPA5*), TLD domain-containing protein 2 (*TLDC2*), D-aminoacyl-tRNA deacylase 1 (*DTD1*), RCC1 and BTB domain-containing protein 2 (*RCBT2*), E3 ubiquitin-protein ligase RNF167 (*RNG1L*), Formin-F (*FH4*), probable GTP-binding protein EngB (*ENGB*), DEAD-box ATP-dependent RNA helicase 17 (*RH17*), and methionine adenosyltransferase 2 subunit beta (*MAT2B*).

**Table 1.** Results of codeml and aBSREL analyses of genes. NA: not annotated.

| Gene name | *S. mic* pepID | codeml |  | aBSREL |  |
| --- | --- | --- | --- | --- | --- |
|  |  | dN/dS | qvalues | dN/dS | qvalues |
| *ABCA5* | Smic12622 | 594.61 | 8.60E-03 | >1000 | 1.94E-02 |
| *ABCD3* | Smic27653 | 26.52 | 2.03E-04 | 12.51 | 6.94E-03 |
| *ARGA* | Smic28624 | 498.45 | 4.03E-03 | 688.52 | 6.51E-03 |
| *ATG26* | Smic28227 | 109.16 | 1.37E-02 | >1000 | 4.26E-02 |
| *CDPK2* | Smic18963 | 238.94 | 1.51E-02 | 899.10 | 4.49E-02 |
| *DTD1* | Smic22294 | 998.94 | 1.37E-02 | >1000 | 2.96E-02 |
| *ENGB* | Smic37911 | 15.80 | 6.85E-04 | 10.30 | 6.51E-03 |
| *FH4* | Smic38689 | 144.87 | 0.00E+00 | 263.34 | 2.08E-02 |
| *MAT2B* | Smic31557 | 7.37 | 1.75E-02 | 13.59 | 0.00E+00 |
| *PPA5* | Smic25730 | 81.79 | 3.86E-04 | >1000 | 1.21E-02 |
| *RCBT2* | Smic7706 | 860.11 | 2.47E-02 | >1000 | 1.21E-02 |
| *RH17* | Smic31560 | 8.84 | 2.11E-02 | 16.11 | 3.44E-02 |
| *RNG1L* | Smic29656 | 270.23 | 1.35E-02 | >1000 | 2.98E-02 |
| *SAC1* | Smic22895 | 999.00 | 3.86E-02 | >1000 | 4.96E-02 |
| *SCNA* | Smic24527 | 193.26 | 4.33E-03 | 640.03 | 3.78E-02 |
| *SSY1* | Smic36738 | 19.09 | 3.32E-03 | 14.68 | 4.92E-02 |
| *TLDC2* | Smic33911 | 999.00 | 7.46E-03 | >1000 | 7.29E-03 |
| NA | Smic36824 | 268.17 | 1.47E-04 | 547.56 | 3.12E-02 |
| NA | Smic38300 | 307.80 | 2.72E-02 | >1000 | 4.41E-02 |
| NA | Smic21495 | 35.54 | 0.00E+00 | 15.66 | 0.00E+00 |
| NA | Smic21475 | 998.99 | 4.67E-04 | 50.34 | 1.93E-02 |
| NA | Smic41910 | 46.26 | 2.03E-04 | 60.02 | 6.51E-03 |
| NA | Smic39357 | 291.35 | 9.49E-04 | 70.54 | 3.12E-02 |
| NA | Smic30582 | 11.47 | 2.57E-02 | 34.61 | 2.63E-02 |
| NA | Smic38876 | 38.35 | 2.94E-03 | 28.75 | 3.99E-02 |
| NA | Smic30203 | 113.95 | 8.58E-03 | >1000 | 3.12E-02 |
| NA | Smic13905 | 290.46 | 2.03E-04 | >1000 | 1.92E-02 |
| NA | Smic24546 | 999.00 | 6.40E-03 | 16.82 | 3.99E-02 |
| NA | Smic32969 | 178.93 | 2.55E-03 | 880.15 | 0.00E+00 |
| NA | Smic9611 | 121.16 | 1.47E-04 | 22.33 | 0.00E+00 |
| NA | Smic43774 | 947.63 | 2.66E-03 | 879.51 | 4.41E-02 |
| NA | Smic33697 | 999.00 | 1.43E-02 | 19.72 | 3.99E-02 |
| NA | Smic30844 | 147.72 | 5.48E-03 | 14.36 | 4.26E-02 |
| NA | Smic22365 | 810.47 | 1.35E-02 | >1000 | 2.89E-02 |
| NA | Smic35783 | 31.12 | 2.03E-04 | >1000 | 0.00E+00 |

### Positively selected sites in soluble starch synthase genes

Among the 35 candidate genes, *SSY1* was particularly notable because sugar metabolism (e.g., starch metabolism) is known to be one of the functions highly relevant to symbiotic relationships (Davy et al. 2012; Gordon & Leggat 2010; Ishii et al. 2019; Maor-Landaw et al. 2020; Mashini et al. 2022; Sproles et al. 2018). Furthermore, in *Crypthecodinium cohnii*, a model dinoflagellate for starch synthesis, mutants deficient in soluble starch synthase activity showed reduced starch content, shortened starch chain length, and reduced starch granule size (Dauvillée et al. 2009). To gain insight into the positive selection of a gene relevant to sugar metabolism, we focused on *SSY1*. From the posterior probability (>0.5) obtained using the Bayes empirical Bayes (BEB) method in the PAML package, 12 amino acid residues were identified as positively selected sites in SSY1 (Table 2). The three-dimensional conformation of *S. mic* SSY1 estimated using the Alphafold2 database (UniProt ID: A0A1Q9CI39) indicated that 8 out of the 12 amino acid residues were located in local structures with very low confidence, of which pLDDTs were smaller than 50. Among the remaining residues, two were located within the protein domain, while the other two were located outside the protein domain. Low confidence local structures were regarded as intrinsically disordered regions, where seven of all positive selected sites were estimated to be located. Compared with other genera, the aforementioned 12 amino acids of symbiotic *Symbiodinium* spp. were not conserved in SSY1 of the symbiotic Symbiodiniaceae species (Figure S1).

**Table 2.** Pfam domains, posterior probability of BEB analysis, pLDDT score, and disorder detection results of selection sites for *SSY1*.

| Gene | Pfam domain | Pfam Short name | selected site | Amino acids of Symbiosis species | Amino acids of free-living species | BEB probabilities<br>Bold>0.95 | pLDDTscore<br>Bold<50 | disorder<br>S: Structured<br>D: Disordered | disorder subtype |
| --- | --- | --- | --- | --- | --- | --- | --- | --- | --- |
| SSY1 | - |  | 11 | L | T | **0.955** | **36.81** | S |  |
|  | PF00686 | CBM_20 | 59 | L | T | **0.952** | 84.25 | S |  |
|  | - |  | 104 | C | T | **0.98** | 90.06 | S |  |
|  | - |  | 135 | L | K | 0.802 | **31.09** | D |  |
|  | - |  | 135 | L | E | 0.802 | **31.09** | D |  |
|  | - |  | 149 | Q | A | **0.964** | **38.59** | D | Polyampholyte |
|  | - |  | 207 | A | S | **0.967** | **30.20** | D | Low complexity |
|  | - |  | 240 | E | P | **0.973** | **26.55** | D |  |
|  | - |  | 241 | A | K | 0.759 | **25.27** | D |  |
|  | - |  | 269 | T | G | 0.545 | **28.23** | D | Low complexity |
|  | - |  | 269 | T | D | 0.545 | **28.23** | D | Low complexity |
|  | - |  | 300 | L | E | **0.967** | **23.17** | D |  |
|  | PF08323 | Glyco_transf_5 | 564 | R | L | 0.569 | 92.00 | S |  |
|  | - |  | 858 | H | C | **0.962** | 77.81 | S |  |

### SSY1 gene family expansion in dinoflagellates

Tandem duplication of exons and genes is commonly found in dinoflagellate genomes (Bachvaroff & Place 2008; Stephens et al. 2020), and it has been suggested as an adaptive mechanism for enhancing biological functions (González-Pech et al. 2021a). Consistent with this, when *SSY1* genes of clades C-F were searched, multiple genes with high homology (E-value 1e-5 in BLASTP) were detected in each genome. To clarify the molecular evolution of the SSY1 gene family, which has undergone positive selection in *Symbiodinium* spp., a phylogenetic analysis of broadly sampled SSY1 was performed. All sequences from the family Symbiodiniaceae (order Suessiales) and other dinoflagellates were reconstructed as a sister monophyletic clade to Cryptophyceae (Figure 2). Within the monophyletic clades of Suessiales, an SSY1 orthogroup containing a positively selected gene product and four additional SSY1 orthogroups of *Symbiodinium* were identified in the phylogenetic tree (Figure 2, Supplementary data 1). Hereafter, they are referred to as orthogroups 1-5, of which Orthogroup 5 is SSY1 with positive selection. Orthogroup 1 included two subgroups (called Orthogroup 1.1 and Orthogroup 1.2). No genes classified into Orthogroup 2 were detected in the CDS datasets of *S. nec*, *S. lin*, *S. nat,* and *S. pil* (Figure S2). No genes classified into Orthogroup 3 were detected in the CDS datasets of *S. mic*04 and *S. pil* (Figure S2). Each monophyletic orthogroup was also closely related to SSY1 homologs of other Symbiodiniaceae families (Figure S2) and various dinoflagellate species, including deeper branching species, such as the genera *Amphidinium* and *Karlodinium* (Janouškovec et al. 2017).

**Fig. 2.**
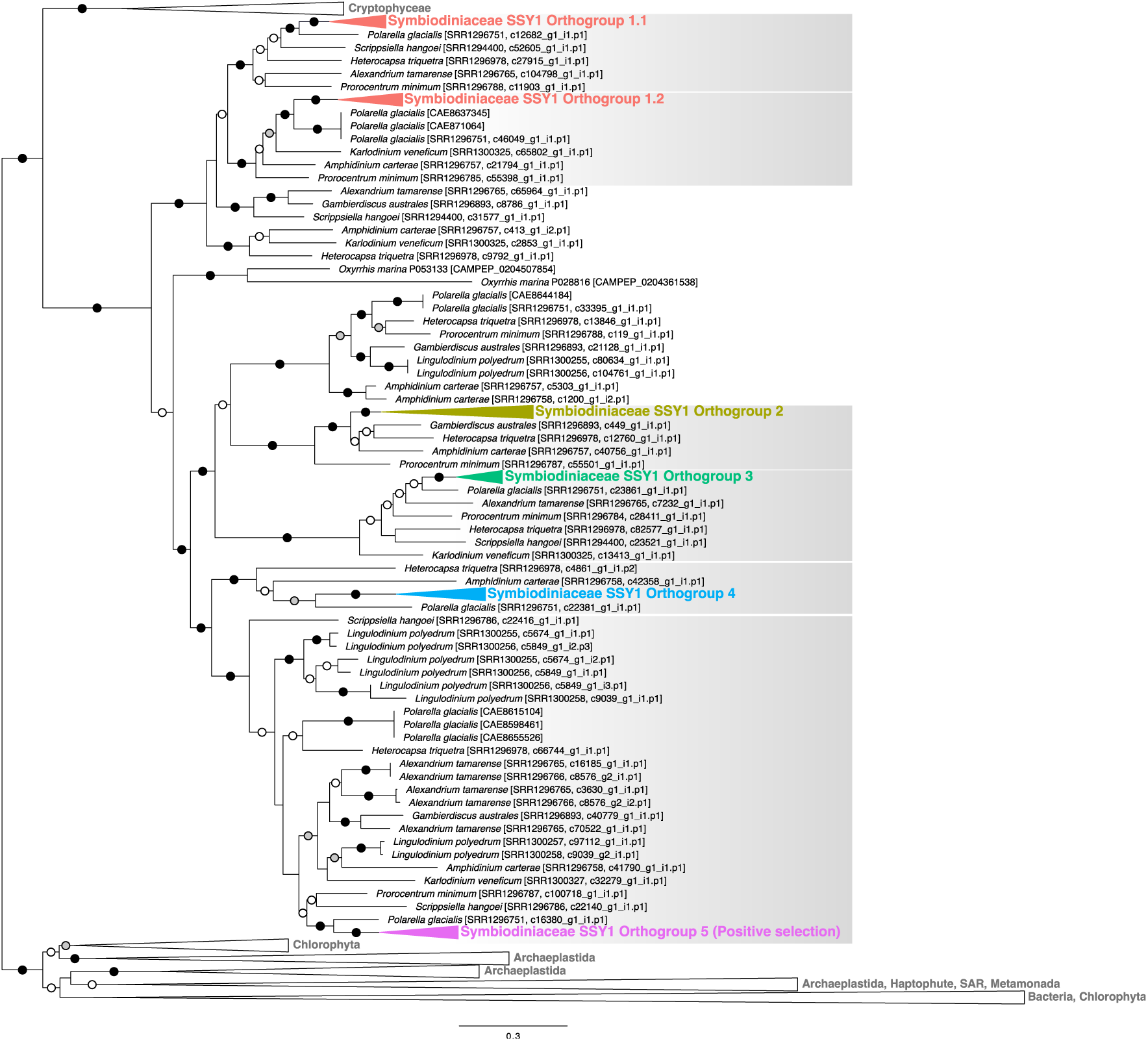
Maximum likelihood tree of SSY1 protein sequences. Simplified tree representing relationships among five orthogroups in *Symbiodinium*. The OTUs of dinoflagellates were labeled with species names. Dinoflagellate SSY1 Orthogroups are shaded in gray. Closed, open gray, and open white circles indicate branches supported with ≥95%, ≥80%, and ≥50% ultrafast bootstrap support, respectively.

To determine whether the positively selected *SSY1* (Orthogroup 5) was transcriptionally active in *Symbiodinium* spp., transcriptome data were evaluated with transcripts per million (TPMs) using publicly available data. Although the TPM was 0 for gene16088 (Orthogroup 1) of *S. tri*, genes of Orthogroup 5 were transcribed as abundantly as most of the other orthogroup gene copies in all the strains investigated in this study (Figure S3).

### Intracellular starch content under nitrogen-deficient and low pH conditions

Given the aforementioned positive selection in *SSY1*, we surveyed the phenotypes with a particular focus on starch in both symbiotic and free-living species. First, we evaluated the molecular characteristics of starch granules in free-living (*S. pil* and *S. nat*) and symbiotic species (*S. tri* and *S. mic*) by measuring the intracellular starch chain lengths and starch granule sizes (Figure S4). When cells cultured in the f/2 medium under typical conditions were analyzed, the distribution of the degree of polymerization (DP), which represents starch chain length, showed peaks of AREA (%) at DP10. The AREA (%) of each species at DP10 was approximately 9-11% for all species analyzed (Figure S4A). The median, maximum, and minimum starch particle sizes measured under the same conditions as the analysis of starch chain length were approximately 0.08-0.09, 0.15-0.19, and 0.03-0.05 μm, respectively. Kruskal–Wallis one-way analysis of variance yielded a *p*-value of 0.7126 (Figure S4B). No significant differences in the chain length or starch granule size were observed among the four species (Figure S4A, B). Based on the fact that starch chain length varies depending on the type of starch synthase (Tetlow & Bertoft 2020), the starch chain lengths of all the four species were comparable to those of semi-amylopectin (also called floridean starch) found in Cryptophyceae and red algae (Nakamura et al. 2005).

Subsequently, we measured the starch content in the cells of free-living (*S. pil* and *S. nat*) and symbiotic species (*S. tri* and *S. mic*) cultured in the f/2 medium. Intriguingly, the intracellular starch content in free-living species was apparently higher than that in symbiotic species. This difference may be due to preferred growth conditions. Thus, we prepared three different media, low pH f/2, nitrogen-deficient f/2, and low pH and nitrogen-deficient f/2 (referred to as f2L, f2N, and f2NL, respectively) (Figure 3), to focus on starch accumulation under actual conditions in the habitat within hosts with low pH and nitrogen deficiency. The average and standard deviation of starch content (µg/9.0×10^8^ cells) under f2, f2L, f2N, and f2NL conditions was 18.48±1.49, 16.08±1.41, 25.07±4.69, and 23.67±2.66 for *S. mic;* 16.97±1.04, 12.98±3.27, 25.49±5.18, and 30.39±2.58 for *S. tri;* 45.07±11.69, 18.05±4.58, 137.44±15.77, and 73.58±20.24 for *S. nat*; and 36.67±11.76, 18.87±2.21, 160.48±23.50, and 66.34±7.00 for *S. pil*. This demonstrated that the starch content of cells cultured in the f/2L medium was reduced by 51% in *S. pil*, 40% in *S. nat*, 76% in *S. tri*, and 87% in *S. mic* compared with that of cells cultured in the f/2 medium, and the reduction was greater in free-living species. The starch content of cells cultured in the f/2N medium increased by 438% in *S. pil*, 305% in *S. nat*, 150% in *S. tri*, and 136% in *S. mic* compared with that of cells cultured in the f/2 medium, and the increase was greater in free-living species. The starch content of cells cultured in the f/2NL medium was 41% in *S. pil*, 53% in *S. nat*, 119% in *S. tri*, and 94% in *S. mic* compared with that of cells cultured in the f/2N medium. The starch content of cells cultured in the f/2NL medium was 319% in *S. pil*, 408% in *S. nat*, 234% in *S. tri*, and 147% in *S. mic* compared with that of cells cultured in the f/2L medium. Overall, starch content fluctuated greatly in response to environmental changes in free-living species, but less substantially in symbiotic species under acidic and nitrogen-deficient conditions, similar to those of the symbiont habitats.

**Fig. 3.**
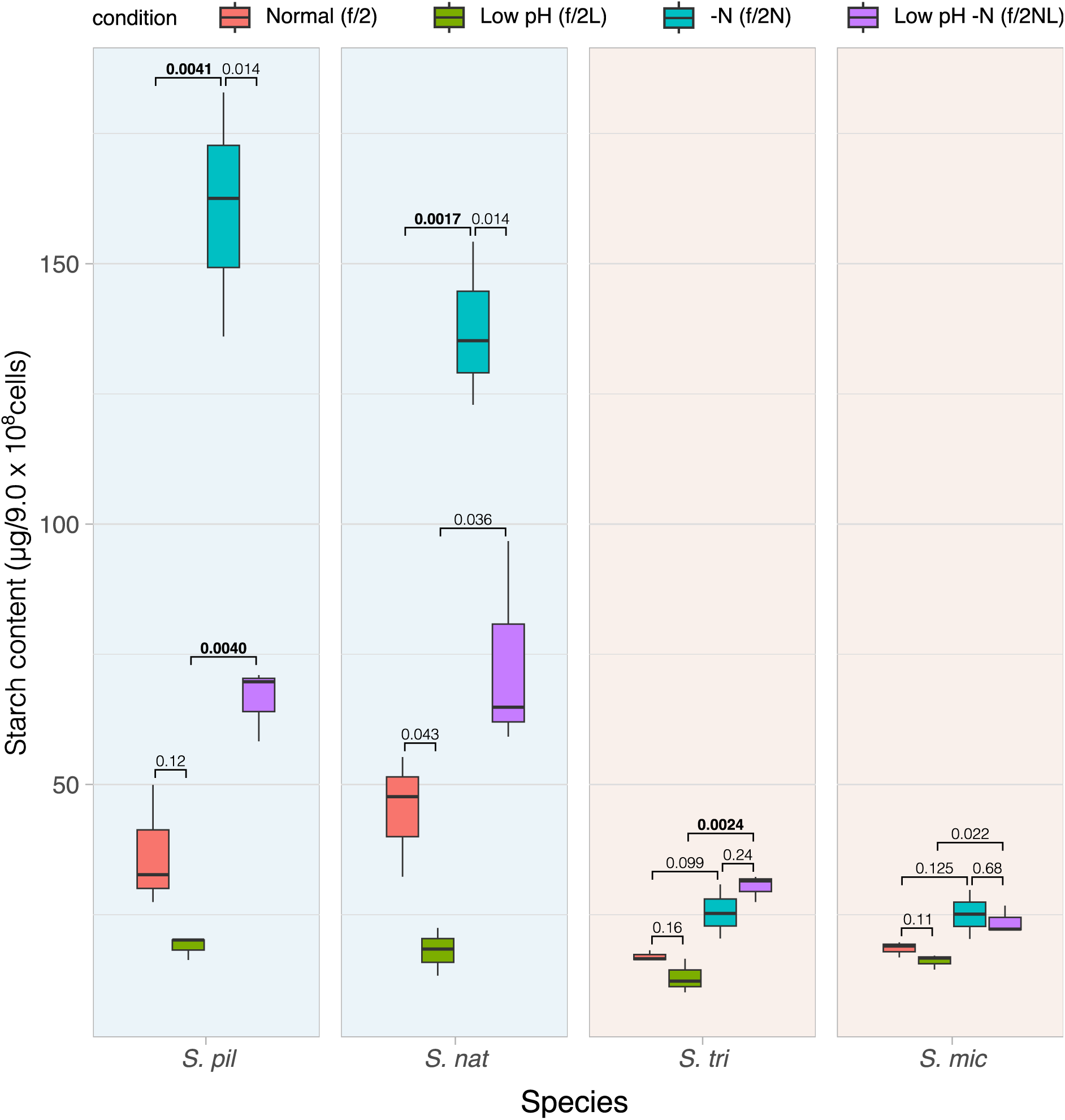
Quantification of starch in cells under different culture conditions. Starch content under different pH and nitrate conditions (n=3 per treatment) using four different media is shown: normal (f2), low pH (f2L), nitrogen-deficient (f2N), and low pH and nitrogen-deficient (f2NL). The numbers in the plots represent *p*-values, with statistically significant differences highlighted in bold (Welch’s two-sample t-test, Bonferroni-corrected *p*<0.005).

## Discussion

A symbiotic lifestyle has been proposed to have emerged during the evolution of Symbiodiniaceae (Gómez 2012; González-Pech et al. 2017; Aranda et al. 2016; Liu et al. 2018). Although some members possess symbiotic lifestyles and others are free-living within the family, even within the genus *Symbiodinium*, evolutionary transitions at the molecular level during free-living and symbiotic lifestyle diversification remain unknown. Here, we present single-gene-level comparative molecular evolution analyses between symbiotic and free-living species of *Symbiodinium*, which allowed us to gain deeper insights into the genetic basis of lifestyle diversification in two largely distinct ways: symbiotic and free-living (Figure 1). Even within the symbiotic species, *S. mic, S. tri, S. lin, and S. nec* were isolated from different hosts such as various species of stony corals, octocorals, or anemones (Aranda et al. 2016; González-Pech et al. 2021a), and especially *S. nec* was found to be opportunistic (González-Pech et al. 2021a; Quigley et al. 2017). Given the host-symbiont relationships varying among species/strains, it is highly likely that each species/strain has evolutionarily adapted to the distinct symbiotic lives with differences in host and stability after their species diversification. Therefore, all the symbiotic species would not entirely share the genetic backbone involved in the symbiotic lifestyles within the genus Symbiodinium. Thus, the 35 positively selected genes detected commonly by two independent *in silico* analyses (Table 1) might include genes of which encoded functions support the principle phenotypes that are common for all the distinct symbiotic lives within the genus Symbiodinium. Of these, 18 genes were not functionally annotated, probably because over 90% of the core genes unique to the genus *Symbiodinium* are “dark matter genes” of unknown function (González-Pech et al. 2021a).

The soluble starch synthase gene encoding *SSY1* was positively selected during lifestyle diversification in *Symbiodinium* (Table 1, Figure 1). Interestingly, most of the positively selected amino acid sites in SSY1 of *Symbiodinium* were found in the predicted intrinsically disordered regions outside the catalytic domains (Table 2). In general, intrinsically disordered regions are characteristic of proteins that are condensed into phase-separated, non-membrane-bound organelles and are involved in many molecular processes, including transcription, translation, and signal transduction (Holehouse & Kragelund 2024). In the model plant *Arabidopsis thaliana*, many starch synthesis proteins also contain intrinsically disordered regions, and liquid-liquid phase separation may be involved in starch synthesis (Bürgy et al. 2021). Given that most amino acid residues at the positively selected were not correlated with the symbiotic/free-living lifestyles in other genera of Symbiodiniaceae (Figure S1), these positive selections likely affected the intra-genus-level evolutionary transitions.

In land plants, starch synthesis-related genes are functionally diversified by gene duplication and work in cooperation, leading to the present form of complex starch synthesis (Cheng et al. 2012; Nougué et al. 2014). We found phylogenetic evidence of gene family expansion and diversification of *SSY1* genes in the common ancestor of the class Dinophyceae, followed by differential loss of either gene copy in certain species/strains (Figure 2). This suggests that multiple copies of *SSY1* generated by gene family expansion may have allowed the accumulation of non-hazardous mutations, followed by natural selection and functional diversification in *Symbiodinium*, as is known for land plants (Cheng et al. 2012; Nougué et al. 2014).

Other genes positively selected during lifestyle diversification were categorized into those possibly involved in metabolic pathways (e.g., *ABCA5, ABCD3, ARGA,* and *ATG26*), extracellular signal sensing (e.g., *SAC1, CDPK2, SCNA*, and *PPA5*), and basic cellular growth and cell cycle (e.g., *TLDC2, DTD1, RCBT2, RNG1L, FH4, ENGB, RH17*, and *MAT2B*). These genes may be involved in symbiosis-related features, such as metabolite exchange between the symbiont and host (Davy et al. 2012), distinct environmental conditions (Armstrong et al. 2018; Barott et al. 2015; Davy et al. 2012; Xiang et al. 2020), and regulation of cell growth in symbiont habitats within the host (Rädecker et al. 2021; Xiang et al. 2020). Notably, *ABCA5* encodes a cholesterol exporter protein that functions in the plasma and lysosomal membranes of mammals (Szakacs & Abele 2020). In the symbiotic relationship between cnidarian hosts and Symbiodiniaceae dinoflagellates, cholesterol and other sterols are essential molecules because cnidarians cannot synthesize them (Baumgarten et al. 2015; Hambleton et al. 2019). Previous studies have proposed sterol secretion by symbionts (Hambleton et al. 2019; Lu et al. 2020), suggesting that sterols are likely to be actively transferred from symbionts via uncharacterized transporters in the “symbiosome” membrane surrounding the Symbiodiniaceae habitat or via unknown pathways. Another example is *ATG26*, which encodes an enzyme that converts sterols, such as cholesterol, to sterol glycosides (Chen et al. 2018a; Warnecke et al. 1999), the latter of which regulates the vertebrate immune system (Murakami-Murofushi & Ohta 1989; Shimamura 2020). Because sterol glycosides have been detected in various microalgae, including dinoflagellates (Yu et al. 2018), *Symbiodinium* is also likely capable of synthesizing sterol glycosides. Interestingly, the host immune system is suppressed when symbionts reside within the host cells, resulting in a stable symbiotic relationship without releasing the symbionts from the host cells (Jacobovitz et al. 2021). However, the entire metabolic pathways, such as starch and sterols, of Symbiodiniaceae remain unclear, and thus, the evolutionary and functional implications for positive selection in the aforementioned positively selected genes remain uncertain based only on *in silico* analysis.

Despite the wealth of comparative genomics data, phenotypic analyses of Symbiodiniaceae species are extremely limited. We found evidence of phenotypic differences in starch accumulation related to lifestyle diversification; starch synthesis/accumulation of symbiotic species was more robust against environmental changes (i.e., pH and nitrogen) than free-living species of *Symbiodinium* (Figure 3). Because the symbiont habitats in the hosts are known to have low pH and nitrogen-deficient conditions, the stability of the carbon metabolite content might have been advantageous in maintaining symbiotic relationships. Although our study did not directly verify the causality among the detected mutations in *SSY1*, variations in starch phenotypes, and lifestyles of *Symbiodinium* spp., it is worth noting that the presence or absence of these characteristics was consistently divided by the foreground branch in the genus *Symbiodinium* (Figure 1). Whether these factors are causally related remains to be investigated in future studies.

Our study identified positively selected genes, including carbon metabolism-related and “dark matter” genes and phenotypic differentiation associated with symbiotic/free-living lifestyle diversification. Although it remains unclear whether the last common ancestor of Symbiodiniaceae was symbiotic or free-living (González-Pech et al. 2021a), the symbiotic species might have undergone independent natural selection after diverging into distinct genera/clades, as exemplified in *SSY1* of *Symbiodinium* (Figure S1). A broader-scale molecular evolution study focusing on other genera of Symbiodiniaceae would help us understand the principal genetic basis linked to the functional changes involved in symbiotic and free-living lifestyle diversification in dinoflagellate-cnidaria symbiotic ecosystems.

## Materials and methods

### Acquisition of CDS datasets

Positively selected genes were detected in the complete set of CDSs in the genome annotations of six *Symbiodinium* species with different lifestyles (symbiotic and free-living). We used CDSs from four symbiotic species, *S. mic*, *S. nec*, *S. lin*, and *S. tri*, and two free-living species, *S. nat* and *S. pil*. We used seven CDS datasets: two CDS datasets of *S. mic* from the strains CCMP2467 (https://phycocosm.jgi.doe.gov/Symmic1/Symmic1.home.html) and 04-503 SCI.03 (Aranda et al. 2016; González-Pech et al. 2021a; González-Pech et al. 2021b), and one CDS dataset each of the other five species (González-Pech et al. 2021a; González-Pech et al. 2021b).

### Ortholog detection and multiple alignments

The CDSs were translated into protein sequences using the TransDecoder (v5.5.0) program (Haas et al. 2013). First, we translated ORFs of sufficient length (minimum protein length: 100 aa) within the CDSs into protein sequences using the TransDecoder program. The detected proteins were subjected to a protein domain search in the Pfam database using hmmer (v3.3.2) (Finn et al. 2011) and a homology search in the Uniprot database using BLASTP (v2.10.1). The results of the homology search in the UniProt database were used to annotate genes. Optimal protein sequences and their corresponding ORFs were obtained based on the results obtained using hmmer and BLASTP. Homologous protein pairs among the strains were estimated using OrthoFinder (v2.5.4) (Emms & Kelly 2019). Genetic trees were constructed for all gene sets identified as single-copy orthogroups. The species tree was constructed using OrthoFineder, and the tree topology was confirmed to be the same as that in previous studies (González-Pech et al. 2021a; LaJeunesse et al. 2018). Genes whose tree topologies matched those of the species tree were used for the subsequent analyses. The gene trees of single-copy orthologs were constructed as follows. The orthologous ORFs from multiple species were aligned together with a codon model using the PRANK (v.170427) multiple alignment program without a user-specified initial guide tree (Löytynoja & Goldman 2005). The aligned sequences were trimmed using TrimAL (v1.4.1) (Capella-Gutiérrez et al. 2009) with an 80% gap threshold (option: -gt 0.8) and then subjected to phylogenetic tree reconstruction using IQ-TREE (1.6.12) (Nguyen et al. 2015) (options: -alrt 1000 -bb 1000), with the model that ModelFinder selected as the best model by likelihood comparison based on the Bayesian information criterion in the default setting. Consistency between the species and gene trees was confirmed by ete-comparison using ETE3 (v3.1.1) (Huerta-Cepas et al. 2016).

### Detection of positive selection

Positive selection was detected for each gene using the aforementioned alignments and constructed trees by two software packages that implemented the branch site model: codeml from the PAML package (Yang 2007), which was used with ete-evol from ETE3 (Huerta-Cepas et al. 2016), and adaptive Branch-Site Random Effects Likelihood (aBSREL) in the HyPhy package (Smith et al. 2015). The detection of genes that underwent positive selection at the branch leading to the exclusive common ancestor of *S. mic, S. nec*, *S. lin,* and *S. tri* is hereafter called the foreground branch (Figure 1). For codeml, two models were fit and tested based on the null and alternative hypotheses of sequence evolution. The null hypothesis assumed that the gene evolved neutrally and the value of ω (i.e., dN/dS) equaled 1 on all the branches (Model A1), whereas the alternative hypothesis assumed that the ω value was >1 on the foreground branch (Model A). The same analyses were conducted using aBSREL implemented in the HyPhy package (Smith et al. 2015). The *q*-values were calculated from a collection of *p*-values for each analysis using codeml and aBSREL, in accordance with the method described by Storey (2002). If the q-value obtained from codeml or aBSREL analysis for a given gene was less than 0.05, it was regarded as a candidate gene that had undergone positive selection at the foreground branch. We used the results of the BEB method obtained using the branch-site model in codeml to check the positively selected amino acid sites.

### Protein structure and phylogenetic analysis of positively selected genes

The tertiary structure of SSY1 protein of *S. mic* was obtained from the AlphaFold Protein Structure Database (Jumper et al. 2021). The intrinsically disordered region of SSY1 protein was detected using MobiDB-lite (Necci et al. 2021). Other sequences used for phylogenetic analyses of *SSY1* were collected by a BLASTP search against the GenBank nr database (as of January 2023); The Comparative Set database of EUKPROT v3; protein sequences of seven *Symbiodinium* strains; and protein sequences of Symbiodiniaceae clades E (Submitted GenBank assembly; GCA_963377175.1), A (Shoguchi et al. 2018), D (Shoguchi 2021), B (Shoguchi et al. 2013), F (Lin et al. 2015; Voolstra et al. 2015), and C (Shoguchi et al. 2018; Voolstra et al. 2015). SSY1 of the ten selected dinoflagellates (*Alexandrium tamarense*, *Amphidinium carterae*, *Gambierdiscus australes*, *Heterocapsa triquetra*, *Karenia brevis*, *Karlodinium veneficum*, *Lingulodinium polyedrum*, *Polarella glacialis*, *Prorocentrum minimum,* and *Scrippsiella hangoei*) was obtained by a TBLASTN search against the Marine Microbial Eukaryote Transcriptome Sequencing Project (MMETSP) transcriptome assembly data (Keeling et al. 2014) using the protein sequences of seven *Symbiodinium* strains as queries. Nucleotide sequences obtained from the transcriptome data were converted to amino acid sequences using TransDecoder in the same manner as for ortholog detection. Almost the same protein sequences of SSY1 Symbiodiniaceae clade E and ten species from the MMETSP transcriptome data were redundantly found and clustered with CD-HIT (v4.8.1) (Li & Godzik 2006) under the 95% sequence identity criterion for each species (option: -c 0.95), and only one representative sequence in each cluster was used for the subsequent analyses. The generated datasets were used for multiple alignments and phylogenetic analyses as previously described (Maruyama et al. 2011), with the following modifications: automatically collected homologous sequences were manually curated and selected for multiple alignments, and IQ-TREE was used to reconstruct phylogenetic trees using the Q.pfam+G4 model, which was selected by ModelFinder as the best model by likelihood comparison based on the Bayesian information criterion (Kalyaanamoorthy et al. 2017; Nguyen et al. 2015). Support values were calculated using the Shimodaira–Hasegawa-like approximate likelihood ratio test (Guindon et al. 2010) and 1000 times ultrafast bootstrap approximation (Hoang et al. 2018).

### Expression levels of SSY1 orthogroup genes

Transcriptome data of *S. mic* (SRR3337493, SRR3337494, and SRR3337498), *S. mic*04 (ERR3711255), *S. lin* (ERR3711290), *S. tri* (ERR4501159), *S. nat* (ERR4501197), and *S. pil* (ERR3711300) were downloaded from the NCBI database. All reads of the fastq files were filtered using fastp (Chen et al. 2018b) (with the options -q 30 and -u 30), and paired output reads were used for the subsequent analyses. The reads from each fastq file were mapped onto each CDS data using STAR 2.7 (Dobin et al. 2013) with default settings. TPMs were calculated using RSEM1.3.3 (Li & Dewey 2011). The TPMs of *S. mic* were calculated as the average TPM from the three transcriptome datasets.

### Strains, culture conditions, and axenic culture generation

*S. mic* (CCMP2467), S*. nat* (CCMP2548), *S. tri* (CCMP2592), and *S. pil* (CCMP2461) were purchased from the National Center for Marine Algae and Microbiota at the Bigelow Laboratory for Ocean Sciences, Maine. *Symbiodinium* strains were maintained according to a previous study (Ishii et al. 2018), with a different medium. Stock cultures were incubated at 26 °C in the f/2 medium made with 36 g/L of Daigo’s Artificial Seawater SP for Marine Microalgae Medium (Shiotani M. S. co., Japan), 20 ml/L of Guillard’s (F/2) Marine Water Enrichment Solution (Sigma-Aldrich, Merck Millipore), and PSN (Gibco, Thermo Fisher Scientific, MA), with final concentrations of penicillin, streptomycin, and neomycin at 0.005, 0.005, and 0.01 mg/mL, respectively. Light was provided at an irradiance of approximately 100 µmol photons/m^2^/s in 12 h light:12 h dark cycles. Axenic cultures were generated as described previously (Costa et al. 2019). Axenic cultures without PSN were cultivated for the subsequent experiments. For the low pH experiments, the pH of the medium was adjusted to 5.5 using HCl. For nitrogen-deficient conditions, the f/2 medium (Guillard 1975) was prepared without NaNO_3_ and Na_2_SiO_3_.

### Quantification of soluble starch

Axenic cultures were inoculated in fresh “normal (f/2),” “low pH (f/2L),” “nitrogen-deficient (f/2N),” or “low pH and nitrogen -deficient (f/2NL)” f/2 media and then incubated for 3 days under the aforementioned temperature and light conditions. Cells with initial densities of 4×10^7^ cells/mL were cultured in 10 ml medium in T25 culture flasks (Cell Cultivation Flask [Vent Cap], AS ONE, Japan). As starch accumulation in algae fluctuates diurnally in association with photosynthesis (Ral et al. 2006), cells were consistently collected 7 h after the beginning of the light period. Cells were collected by centrifugation of 9 ml of culture at 2000×g for 2 min at room temperature, suspended in 1 ml of 80% EtOH, and stored at 4 °C until measurement. Two hundred and fifty microliters of glass beads (425-600 μm in diameter, Cat# G8772, Sigma-Aldrich, USA) were added to a suspension of 9.0×10^8^ cells, and the cells were disrupted with a bead crusher (μT-12, TAITEC, Japan) for 1 min at 3200 rpm with cooling on ice in sextuplicate. The cell lysates were filtered through a 30 μm syringe Filcon Sterite (Cat#340606, DB, USA) and centrifuged at 10,000×g for 20 min at 15 °C to obtain pellets. The following steps were repeated twice to wash the pellets. The pellets were suspended in 90% EtOH, heated at 60 °C for 5 min and centrifuged at 10,000×g for 2 min, followed by removal of the supernatant. To extract soluble starch, the pellets were resuspended in 400 µl of H_2_O and incubated in boiling water for 5 min. The soluble starch extracts were centrifuged at 10,000×g for 2 min, the supernatant was collected, and the amount of starch was measured. The EnzyChrom Starch Assay Kit (Cat# E2ST-100, BioAssay Systems, USA) and a plate reader (Biotek cytation5, Agilent Technologies, USA) were used for measurement, following the manufacturer’s instructions. The obtained data were analyzed using R.

### Comparison of the chain length distribution profiles and the granule size of starch among the strains

Cells in 20 ml cell cultures in the exponential growth phase were collected by centrifugation. After removing the medium, the cells were resuspended in 1 ml of 80% EtOH. Two hundred and fifty microliters of glass beads (425-600 μm in diameter) and 10 μl of smaller glass beads (≤106 μm in diameter, Cat# G4649, Sigma-Aldrich) were added to the cell suspension, and the cells were disrupted by vortexing for 5 min. The cell lysates were filtered through a 30 μm syringe Filcon Sterite and centrifuged at 10,000×g for 20 min at 15 °C to obtain pellets. The pellets were resuspended in 60% EtOH, vortexed, and centrifuged at 10,000×g for 2 min, followed by removal of the supernatant. The pellets were resuspended in 0.8 ml of 10% EtOH, vortexed, and centrifuged at 10,000×g for 20 min at 15 °C, followed by removal of the supernatant. The pellets were resuspended in 10% EtOH, resulting in a total of 0.9 ml, the crude fraction (CF). The CF was subjected to analysis of chain length distribution profiles (Nakamura et al. 2005) and SEM observations (Shimonaga et al. 2008) in the same manner as described in a previous study. The diameter of the starch granules in the SEM images was estimated using ImageJ (Schneider et al. 2012).

## Data availability

The dataset supporting the results of this study is included in the article and supplementary material files.

## Acknowledgments

We thank Prof. Noriyuki Kioka and Dr. Mito Kuroda for their help in measuring the amount of starch, Yuna Uchida and Asuka Kodaira for their assistance in maintaining the algal cultures, and Daisuke Yamagishi for his help during the early phases of the analysis. The experiments were partially performed using the NIG supercomputer at the ROIS National Institute of Genetics. Computation time was provided by the Super Computer System at the Institute for Chemical Research, Kyoto University. This work was supported in part by JSPS KAKENHI (grant numbers 20K15871, 23KJ2228, and 24K18190 [awarded to Y.I.]; 23K23960, 23H04962, and 24H01462 [awarded to S.M.]; 24K21929 [awarded to R.K.]; and 21H05057 [awarded to T.Y.]). Structural analysis of starch was performed with the help of Starch Technologies Co., Ltd. (https://starchtec.com).

## Conflict of interest

Authors declare that they have no competing interests.

## Notes

### Competing Interest Statement

The authors have declared no competing interest.

## References

Aranda M et al. 2016. Genomes of coral dinoflagellate symbionts highlight evolutionary adaptations conducive to a symbiotic lifestyle. Sci Rep. 6:39734. doi: 10.1038/srep39734.

Armstrong EJ, Roa JN, Stillman JH, Tresguerres M. 2018. Symbiont photosynthesis in giant clams is promoted by V-type H^+^-ATPase from host cells. J Exp Biol. 221. doi: 10.1242/jeb.177220.

Bachvaroff TR, Place AR. 2008. From stop to start: tandem gene arrangement, copy number and trans-splicing sites in the dinoflagellate *Amphidinium carterae*. PLoS One. 3:e2929. doi: 10.1371/journal.pone.0002929.

Baker AC. 2003. Flexibility and specificity in coral-algal symbiosis: diversity, ecology, and biogeography of *Symbiodinium*. Annu Rev Ecol Evol Syst. 34:661–689. doi: 10.1146/annurev.ecolsys.34.011802.132417.

Barott KL, Venn AA, Perez SO, Tambutté S, Tresguerres M. 2015. Coral host cells acidify symbiotic algal microenvironment to promote photosynthesis. Proc Natl Acad Sci U.S.A. 112:607–612. doi: 10.1073/pnas.1413483112.

Baumgarten S et al. 2015. The genome of *Aiptasia*, a sea anemone model for coral symbiosis. Proc Natl Acad Sci U.S.A. 112:11893–11898.

Blackall LL, Wilson B, van Oppen MJH. 2015. Coral—the world’s most diverse symbiotic ecosystem. Mol Ecol. 24:5330–5347. doi: 10.1111/mec.13400.

Bürgy L et al. 2021. Coalescence and directed anisotropic growth of starch granule initials in subdomains of *Arabidopsis thaliana* chloroplasts. Nat Commun. 12:6944. doi: 10.1038/s41467-021-27151-5.

Burriesci MS, Raab TK, Pringle JR. 2012. Evidence that glucose is the major transferred metabolite in dinoflagellate-cnidarian symbiosis. J Exp Biol. 215:3467–3477. doi: 10.1242/jeb.070946.

Capella-Gutiérrez S, Silla-Martínez JM, Gabaldón T. 2009. trimAl: a tool for automated alignment trimming in large-scale phylogenetic analyses. Bioinformatics. 25:1972–1973. doi: 10.1093/bioinformatics/btp348.

Chen Liuqing, Zhang Y, Feng Y. 2018a. Structural dissection of sterol glycosyltransferase UGT51 from *Saccharomyces cerevisiae* for substrate specificity. J Struct Biol. 204:371–379. doi: 10.1016/j.jsb.2018.11.001.

Chen Shifu, Zhou Y, Chen Y, Gu J. 2018b. fastp: an ultra-fast all-in-one FASTQ preprocessor. Bioinformatics. 34:i884–i890. doi: 10.1093/bioinformatics/bty560.

Cheng J et al. 2012. Diversification of genes encoding granule-bound starch synthase in monocots and dicots Is marked by multiple genome-wide duplication events. PLoS One. 7:e30088. doi: 10.1371/journal.pone.0030088.

Costa RM, Fidalgo C, Cárdenas A, Frommlet JC, Voolstra C. 2019. Protocol for the generation of axenic/bacteria-depleted Symbiodiniaceae cultures. doi: 10.17504/protocols.io.87khzkw

Cui G et al. Molecular insights into the Darwin paradox of coral reefs from the sea anemone *Aiptasia*. Sci Adv. 9:eadf7108. doi: 10.1126/sciadv.adf7108.

Dauvillée D et al. 2009. Genetic dissection of floridean starch synthesis in the cytosol of the model dinoflagellate *Crypthecodinium cohnii*. Proc Natl Acad Sci U.S.A. 106:21126–21130. doi: 10.1073/pnas.0907424106.

Davy SK, Allemand D, Weis VM. 2012. Cell biology of cnidarian-dinoflagellate symbiosis. Microbiol Mol Biol Rev. 76:229–261. doi: 10.1128/MMBR.05014-11.

Dimijian GG. 2000. Evolving together: the biology of symbiosis, Part 1. Proc (Bayl Univ Med Cent). 13:217a–2226. doi: 10.1080/08998280.2000.11927678.

Dobin A et al. 2013. STAR: ultrafast universal RNA-seq aligner. Bioinformatics. 29:15–21. doi: 10.1093/bioinformatics/bts635.

Emms DM, Kelly S. 2019. OrthoFinder: phylogenetic orthology inference for comparative genomics. Genome Biol. 20:238. doi: 10.1186/s13059-019-1832-y.

Finn RD, Clements J, Eddy SR. 2011. HMMER web server: interactive sequence similarity searching. Nucleic Acids Res. 39:W29–W37. doi: 10.1093/nar/gkr367.

Frankowiak K et al. 2016. Photosymbiosis and the expansion of shallow-water corals. Sci Adv. 2:e1601122. doi: 10.1126/sciadv.1601122.

Gilbert SF, Sapp J, Tauber AI. 2012. A symbiotic view of life: we have never been individuals. Q Rev Biol. 87:325–341. doi: 10.1086/668166.

Gómez F. 2012. A quantitative review of the lifestyle, habitat and trophic diversity of dinoflagellates (Dinoflagellata, Alveolata). Syst Biodivers. 10:267–275. doi: 10.1080/14772000.2012.721021.

González-Pech RA, Stephens TG, et al. 2021a. Comparison of 15 dinoflagellate genomes reveals extensive sequence and structural divergence in family Symbiodiniaceae and genus *Symbiodinium*. BMC Biol. 19:73. doi: 10.1186/s12915-021-00994-6.

González-Pech RA, Ragan MA, Bhattacharya D, Chan CX. 2021b. Genome assemblies and the associated annotations for seven *Symbiodinium* isolates. The University of Queensland. Data Collection. doi: 10.14264/f1b3a11.

González-Pech RA, Ragan MA, Chan CX. 2017. Signatures of adaptation and symbiosis in genomes and transcriptomes of *Symbiodinium*. Sci Rep. 7:15021. doi: 10.1038/s41598-017-15029-w.

Gordon BR, Leggat W. 2010. *Symbiodinium*—invertebrate symbioses and the role of metabolomics. Mar Drugs. 8:2546–2568. doi: 10.3390/md8102546.

Grant AJ, Rémond M, Starke-Peterkovic T, Hinde R. 2006. A cell signal from the coral *Plesiastrea versipora* reduces starch synthesis in its symbiotic alga, Symbiodinium sp. Comp Biochem Physiol A. 144:458–463. doi: 10.1016/j.cbpa.2006.04.012.

Guillard RRL. 1975. Culture of phytoplankton for feeding marine invertebrates. In: Culture of Marine Invertebrate Animals. Springer, Boston, MA pp. 29–60. doi: 10.1007/978-1-4615-8714-9_3.

Guindon S et al. 2010. New algorithms and methods to estimate maximum-likelihood phylogenies: assessing the performance of PhyML 3.0. Syst Biol. 59:307–321. doi: 10.1093/sysbio/syq010.

Haas BJ et al. 2013. *De novo* transcript sequence reconstruction from RNA-seq using the Trinity platform for reference generation and analysis. Nat Protoc. 8:1494–1512. doi: 10.1038/nprot.2013.084.

Hambleton EA et al. 2019. Sterol transfer by atypical cholesterol-binding NPC2 proteins in coral-algal symbiosis. eLife. 8:e43923. doi: 10.7554/eLife.43923.

Hansen G, Daugbjerg N. 2009. *Symbiodinium natans* sp. nov.: a “free-living” dinoflagellate from tenerife (northeast-atlantic ocean)1. J Phycol. 45:251–263. doi: 10.1111/j.1529-8817.2008.00621.x.

Hoang DT, Chernomor O, von Haeseler A, Minh BQ, Vinh LS. 2018. UFBoot2: improving the ultrafast bootstrap approximation. Mol Biol Evol. 35:518–522. doi: 10.1093/molbev/msx281.

Holehouse AS, Kragelund BB. 2024. The molecular basis for cellular function of intrinsically disordered protein regions. Nat Rev Mol Cell Biol. 25:187–211. doi: 10.1038/s41580-023-00673-0.

Huerta-Cepas J, Serra F, Bork P. 2016. ETE 3: Reconstruction, analysis, and visualization of phylogenomic data. Mol Biol Evol. 33:1635–1638. doi: 10.1093/molbev/msw046.

Ishii Y et al. 2023. Environmental pH signals the release of monosaccharides from cell wall in coral symbiotic alga Tessmar-Raible, K & Schuman, MC, editors. eLife. 12:e80628. doi: 10.7554/eLife.80628.

Ishii Y et al. 2019. Global shifts in gene expression profiles accompanied with environmental changes in cnidarian-dinoflagellate endosymbiosis. G3-GENES GENOM GENET. 9:2337–2347. doi: 10.1534/g3.118.201012.

Ishii Y et al. 2018. Isolation of uracil auxotroph mutants of coral symbiont alga for symbiosis studies. Sci Rep. 8:3237. doi: 10.1038/s41598-018-21499-3.

Jacobovitz MR et al. 2021. Dinoflagellate symbionts escape vomocytosis by host cell immune suppression. Nat Microbiol. 1–14. doi: 10.1038/s41564-021-00897-w.

Janouškovec J et al. 2017. Major transitions in dinoflagellate evolution unveiled by phylotranscriptomics. Proc Natl Acad Sci U.S.A. 114:E171–E180. doi: 10.1073/pnas.1614842114.

Jumper J et al. 2021. Highly accurate protein structure prediction with AlphaFold. Nature. 596:583–589. doi: 10.1038/s41586-021-03819-2.

Kalyaanamoorthy S, Minh BQ, Wong TKF, von Haeseler A, Jermiin LS. 2017. ModelFinder: fast model selection for accurate phylogenetic estimates. Nat Methods. 14:587–589. doi: 10.1038/nmeth.4285.

Keeling PJ et al. 2014. The Marine Microbial Eukaryote Transcriptome Sequencing Project (MMETSP): Illuminating the functional diversity of eukaryotic life in the oceans through transcriptome sequencing. PLoS Biol. 12:e1001889. doi: 10.1371/journal.pbio.1001889.

LaJeunesse TC et al. 2018. Systematic revision of Symbiodiniaceae highlights the antiquity and diversity of coral endosymbionts. Curr Biol. 28:2570–2580.e6. doi: 10.1016/j.cub.2018.07.008.

Li B, Dewey CN. 2011. RSEM: accurate transcript quantification from RNA-Seq data with or without a reference genome. BMC Bioinform. 12:323. doi: 10.1186/1471-2105-12-323.

Li W, Godzik A. 2006. Cd-hit: a fast program for clustering and comparing large sets of protein or nucleotide sequences. Bioinformatics. 22:1658–1659. doi: 10.1093/bioinformatics/btl158.

Lin S et al. 2015. The *Symbiodinium kawagutii* genome illuminates dinoflagellate gene expression and coral symbiosis. Science. 350:691–694. doi: 10.1126/science.aad0408.

Liu H et al. 2018. *Symbiodinium* genomes reveal adaptive evolution of functions related to coral-dinoflagellate symbiosis. Commun Biol. 1:1–11. doi: 10.1038/s42003-018-0098-3.

Löytynoja A, Goldman N. 2005. An algorithm for progressive multiple alignment of sequences with insertions. Proc Natl Acad Sci U.S.A. 102:10557–10562. doi: 10.1073/pnas.0409137102.

Lu Y et al. 2020. Clade-specific sterol metabolites in dinoflagellate endosymbionts are associated with coral bleaching in response to environmental cues. mSystems. 5:e00765–20. doi: 10.1128/mSystems.00765-20.

Maor-Landaw K et al. 2023. A candidate transporter allowing symbiotic dinoflagellates to feed their coral hosts. ISME Commun. 3:1–7. doi: 10.1038/s43705-023-00218-8.

Maor-Landaw K, van Oppen MJH, McFadden GI. 2020. Symbiotic lifestyle triggers drastic changes in the gene expression of the algal endosymbiont *Breviolum minutum* (Symbiodiniaceae). Ecol Evol. 10:451–466. doi: 10.1002/ece3.5910.

Maruyama S, Suzaki T, Weber AP, Archibald JM, Nozaki H. 2011. Eukaryote-to-eukaryote gene transfer gives rise to genome mosaicism in euglenids. BMC Evol Biol. 11:105. doi: 10.1186/1471-2148-11-105.

Mashini AG, Oakley CA, Grossman AR, Weis VM, Davy SK. 2022. Immunolocalization of metabolite transporter proteins in a model cnidarian-dinoflagellate symbiosis. Appl Environ Microbiol. 88:e00412–22. doi: 10.1128/aem.00412-22.

McFall-Ngai M et al. 2013. Animals in a bacterial world, a new imperative for the life sciences. Proc Natl Acad Sci U.S.A. 110:3229–3236. doi: 10.1073/pnas.1218525110.

Mies M, Sumida PYG, Radecker N, Voolstra CR. 2017. Marine invertebrate larvae associated with *Symbiodinium*: A mutualism from the start? Front Ecol Evol. doi: 10.3389/fevo.2017.00056.

Murakami-Murofushi K, Ohta J. 1989. Expression of UDP-glucose: poriferasterol glucosyltransferase in the process of differentiation of a true slime mold, *Physarum polycephalum*. Biochim Biophys Acta. 992:412–415. doi: 10.1016/0304-4165(89)90108-6.

Muscatine L. 1990. The role of symbiotic algae in carbon and energy flux in reef corals. Coral Reefs. 25:75–87.

Nakamura Y et al. 2005. Some cyanobacteria synthesize semi-amylopectin type α-polyglucans instead of glycogen. Plant Cell Physiol. 46:539–545. doi: 10.1093/pcp/pci045.

Necci M, Piovesan D, Clementel D, Dosztányi Z, Tosatto SCE. 2021. MobiDB-lite 3.0: fast consensus annotation of intrinsic disorder flavors in proteins. Bioinformatics. 36:5533–5534. doi: 10.1093/bioinformatics/btaa1045.

Nguyen L-T, Schmidt HA, von Haeseler A, Minh BQ. 2015. IQ-TREE: a fast and effective stochastic algorithm for estimating maximum-likelihood phylogenies. Mol Biol Evol. 32:268–274. doi: 10.1093/molbev/msu300.

Norton JH, Shepherd MA, Long HM, Fitt WK. 1992. The zooxanthellal tubular system in the giant clam. Biol Bull. 183:503–506. doi: 10.2307/1542028.

Nougué O, Corbi J, Ball SG, Manicacci D, Tenaillon MI. 2014. Molecular evolution accompanying functional divergence of duplicated genes along the plant starch biosynthesis pathway. BMC Evol Biol. 14:103. doi: 10.1186/1471-2148-14-103.

Quigley KM, Bay LK, Willis BL. 2017. Temperature and water quality-related patterns in sediment-associated *Symbiodinium* communities impact symbiont uptake and fitness of juveniles in the genus *Acropora*. Front Mar Sci. 4:401. doi: 10.3389/fmars.2017.00401

Rädecker N et al. 2021. Heat stress destabilizes symbiotic nutrient cycling in corals. Proc Natl Acad Sci U.S.A. 118:e2022653118. doi: 10.1073/pnas.2022653118.

Ral J-P et al. 2006. Circadian clock regulation of starch metabolism establishes GBSSI as a major contributor to amylopectin synthesis in *Chlamydomonas reinhardtii*. Plant Physiol. 142:305–317. doi: 10.1104/pp.106.081885.

Roth MS. 2014. The engine of the reef: photobiology of the coral-algal symbiosis. Front Microbiol. 22:5:422. doi: 10.3389/fmicb.2014.00422.

Schneider CA, Rasband WS, Eliceiri KW. 2012. NIH Image to ImageJ: 25 years of image analysis. Nat Methods. 9:671–675. doi: 10.1038/nmeth.2089.

Shimamura M. 2020. Structure, metabolism and biological functions of steryl glycosides in mammals. Biochem J. 477:4243–4261. doi: 10.1042/BCJ20200532.

Shimonaga T et al. 2008. Variation in storage α-glucans of the Porphyridiales (Rhodophyta). Plant Cell Physiol. 49:103–116. doi: 10.1093/pcp/pcm172.

Shoguchi E et al. 2013. Draft assembly of the *Symbiodinium minutum* nuclear genome reveals dinoflagellate gene structure. Curr Biol. 23:1399–1408. doi: 10.1016/j.cub.2013.05.062.

Shoguchi E. 2021. Gene clusters for biosynthesis of mycosporine-like amino acids in dinoflagellate nuclear genomes: Possible recent horizontal gene transfer between species of Symbiodiniaceae (Dinophyceae). J Phycol. doi: 10.1111/jpy.13219.

Shoguchi E et al. 2018. Two divergent *Symbiodinium* genomes reveal conservation of a gene cluster for sunscreen biosynthesis and recently lost genes. BMC Genomics. 19:458. doi: 10.1186/s12864-018-4857-9.

Simpson C, Kiessling W, Mewis H, Baron-Szabo RC, Müller J. 2011. Evolutionary diversification of reef corals: a comparison of the molecular and fossil records. Evolution. 65:3274–3284. doi: 10.1111/j.1558-5646.2011.01365.x.

Smith MD et al. 2015. Less is more: an adaptive branch-site random effects model for efficient detection of episodic diversifying selection. Mol Biol Evol. 32:1342–1353. doi: 10.1093/molbev/msv022.

Sproles AE et al. 2018. Phylogenetic characterization of transporter proteins in the cnidarian-dinoflagellate symbiosis. Mol Phylogenet Evol. 120:307–320. doi: 10.1016/j.ympev.2017.12.007.

Stephens TG et al. 2020. Genomes of the dinoflagellate *Polarella glacialis* encode tandemly repeated single-exon genes with adaptive functions. BMC Biol. 18:1–21. doi: 10.1186/s12915-020-00782-8.

Storey JD. 2002. A direct approach to false discovery rates. J R Stat Soc Series B Stat Methodol. 64:479–498. doi: 10.1111/1467-9868.00346.

Szakacs G, Abele R. 2020. An inventory of lysosomal ABC transporters. FEBS Lett. 594:3965–3985. doi: 10.1002/1873-3468.13967.

Tetlow IJ, Bertoft E. 2020. A review of starch biosynthesis in relation to the building block-backbone model. Int J Mol Sci. 21:7011. doi: 10.3390/ijms21197011.

Voolstra C et al. 2015. The ReFuGe 2020 Consortium—using “omics” approaches to explore the adaptability and resilience of coral holobionts to environmental change. Front Mar Sci. 2:68. doi: 10.3389/fmars.2015.00068

Warnecke D et al. 1999. Cloning and functional expression of *UGT* genes encoding sterol glucosyltransferases from *Saccharomyces cerevisiae*, *Candida albicans*, *Pichia pastoris*, and *Dictyostelium discoideum*. J Biol Chem. 274:13048–13059. doi: 10.1074/jbc.274.19.13048.

Weng L-C et al. 2014. Nitrogen deprivation induces lipid droplet accumulation and alters fatty acid metabolism in symbiotic dinoflagellates isolated from *Aiptasia pulchella*. Sci Rep. 4:5777. doi: 10.1038/srep05777.

Xiang T et al. 2020. Symbiont population control by host-symbiont metabolic interaction in Symbiodiniaceae-cnidarian associations. Nat Commun. 11:108. doi: 10.1038/s41467-019-13963-z.

Yang Z. 2007. PAML 4: Phylogenetic analysis by maximum likelihood. Mol Biol Evol. 24:1586–1591. doi: 10.1093/molbev/msm088.

Yellowlees D, Rees TAV, Leggat W. 2008. Metabolic interactions between algal symbionts and invertebrate hosts. Plant Cell Environ. 31:679–694. doi: 10.1111/j.1365-3040.2008.01802.x.

Yorifuji M et al. 2021. Unique environmental Symbiodiniaceae diversity at an isolated island in the northwestern Pacific. Mol Phylogenet Evol. 107158. doi: 10.1016/j.ympev.2021.107158.

Yu S et al. 2018. Characterization of steryl glycosides in marine microalgae by gas chromatography–triple quadrupole mass spectrometry (GC–QQQ-MS). J Sci Food Agric. 98:1574–1583. doi: 10.1002/jsfa.8629.

