## Supplementary data legend for "Positive selection of a starch synthesis gene and phenotypic differentiation of starch accumulation in symbiotic and free-living coral symbiont dinoflagellate species"

**Supplementary figure 1.** Alignment of positively selected amino acids within SSY1, including other Symbiodiniaceae clades.

Sequences of clades C, B, and D used in this study are of symbiotic species, and those of clades F and E are of free-living species. While the amino acids of clades C-F corresponding to positions 11, 207, 240, 241, and 269 in clade A (*Symbiodinium*) could not be confidently identified because of the ambiguously aligned regions, the amino acids of clades C-F corresponding to positions 59, 104, 149, and 564 of *Symbiodinium* were identical to those of free-living species of *Symbiodinium*.

**Supplementary figure 2. Maximum likelihood tree of SSY1 orthogroups in Symbiodiniaceae**

Trees representing relationships inside five orthogroups in Symbiodiniaceae. Closed, open gray, and open white circles indicate branches supported with ≥95%, ≥80%, and ≥50% ultrafast bootstrap support, respectively. OG: Orthogroup, PS: Positive selection

**Supplementary figure 3. Expression profiles of all *SSY1* genes in *Symbiodinium***

TPMs of all *SSY1* genes in the strains used in this analysis. Colors represent genes in different orthogroups; purple indicates positively selected genes in Orthogroup 5 (OG5_PS). In *S. mic*, six *SSY1* genes belonging to five orthogroups were transcribed.

**Supplementary figure 4. Characteristics of starch granules in free-living and symbiotic *Symbiodinium* spp.**

A. Comparison of chain length distribution (DP) in *Symbiodinium* spp. B. Distribution of starch granule size in *Symbiodinium* spp.

**Supplementary data.** Unabbreviated SSY1 tree
