## Supplementary figure 1 for "Positive selection of a starch synthesis gene and phenotypic differentiation of starch accumulation in symbiotic and free-living coral symbiont dinoflagellate species"

| clade | Species | symbiotic or free living | gene ID or genome position | Amino acid position 11 | 59 | 104 | 135 | 149 | 207 |
|---|---|---|---|---|---|---|---|---|---|
| C | Cladocopium goreaui | S | scaffold1774:40552-48657(+) | -------------------- | MALALTTTAD**T**FPTWQSA-PI | WQKFDGNYRV**T**PVAGQVLQAV | LE-VPEEPPV**E**---------- | ------DKAI**A**QQENEERLLK | ---AP-EPEK**A**---------- |
| F | Fugacium kawagutii | F | scaffold3108:27584-39530(+) | -------------------- | MALPLTTSAD**T**FPVWQSA-PI | WQKFEGNYHV**T**PVDGKVLQAL | LSEAP---------------- | ------DKAA**A**KQDHEEVLLK | ---APGEEAK**E**---------- |
| B | Breviolum minutum | S | 001876_t1 | KFQDVLPGSA**L**TLSSALVGI | MALPLTTSAD**T**FPTWQSA-PV | WQKFDGNYRV**T**PVAGQVLQAV | LE-VPED-LT**E**---------- | ------DKAA**A**KHEKEEILLK | ---GPGEAQK**E**---------- |
| D | Durusdinium trenchi | S | g21608.t1 | -------------------- | -------------------- | --------------------- | --------------------- | --------------------- | --------------------- |
| E | Effrenium voratum | F | CAJ1456987 | -------------------- | MALPLTTSAD**T**FPEWQSS-AI | WQNFQGNYRV**T**PVAGKLIQAA | SL--PETVQT**R**AEGKGTKGTQ | ADKAAADKAA**A**KHEKEEILLK | IPGTGTQGTP**S**---------- |
| A | Symbiodinium pilosum | F | gene5801 | --------TA**T**PA------- | MALPMTTSAD**T**FPNWQSD-PV | WQNFQGNYRV**T**PVTGEVLKAA | LG--P-FLPK**K**V--------- | ------GKAR**A**KQEKEEILLK | ---TETEAKE**S**---------- |
| A | Symbiodinium natans | F | gene5799 | --------KA**T**P-------- | MALPLETSKD**T**FPKWRSAKPV | WQSFQGNYRV**T**PVANEVLLAT | LG--P-SGPS**E**---------- | ------DRAA**A**KQEKEEILLK | -----TEPSP**S**HAQDKDVKDV |
| A | Symbiodinium tridacnidorum | S | gene20042 | --------SS**L**T-------- | MALPLQTSAE**L**FPKWQTVKPV | WQNFQGNYRV**C**PVADEVLLAT | LG--PASSTT**L**---------- | ------DRAV**Q**KQEKEEILLK | -----SEPQA**A**---------- |
| A | Symbiodinium linucheae | S | gene10151 | --------SS**L**T-------- | MALPLQTSAE**L**FPKWQTVKPV | WQNFQGNYRV**C**PVADEVLLAT | LG--PASSTT**L**---------- | ------DRAV**Q**KQEKEEILLK | -----SEPQA**A**---------- |
| A | Symbiodinium necroappetens | S | gene13677 | -------GSS**L**T-------- | MALPLQTSAE**L**FPKWQTVKPV | WQNFQGNYRV**C**PVADEVLLAA | LG--PASSTT**L**---------- | ------DRAV**Q**KQEKEEILLK | -----SEPQA**A**---------- |
| A | Symbiodinium microadriaticum 04 | S | gene21048 | -------GSS**L**T-------- | MALPLQTSAE**L**FPKWQTVKPV | WQNFQGNYRV**C**PVADEVLLAA | LG--PASSTT**L**---------- | ------DRAV**Q**KQEKEEILLK | -----SEPQA**A**---------- |
| A | Symbiodinium microadriaticum | S | gene36738 | -------GSS**L**T-------- | MALPLQTSAE**L**FPKWQTVKPV | WQNFQGNYRV**C**PVADEVLLAA | LG--PASSTT**L**---------- | ------DRAV**Q**KQEKEEILLK | -----SEPQA**A**---------- |

| clade | Species | symbiotic or free living | gene ID or genome position | Amino acid position 240, 241 | 269 | 300 | 564 | 858 |
|---|---|---|---|---|---|---|---|---|
| C | Cladocopium goreaui | S | scaffold1774:40552-48657(+) | --------EG**EK**-----TEGT- | -VKEVN--ES**T**AEDADA-QAE | VTQAPMD-DE**D**EAVEYEVPLK | IDSKAAHDLV**L**GNCINLSKGA | ERQIDIVLYE**P**---------A |
| F | Fugacium kawagutii | F | scaffold3108:27584-39530(+) | ---------------KPPEST- | -PSKEP--EA**T**-------EPT | VSQQPMEEEE**G**EAVEYEMPLK | -DFSAARDLV**L**GNCINLSKGA | --------------------- |
| B | Breviolum minutum | S | 001876_t1 | --------KV**EK**P--KEPEST- | --PKE---PE**T**AVDA------- | VTQAPLV-DE**D**EAEEYEVPLK | LDDKAAHDLV**L**GNCINLSKGA | ERQIDIVLYE**C**---------A |
| D | Durusdinium trenchi | S | g21608.t1 | ---------------------- | --------------------- | --------------------- | IDDKAAGDLV**L**GNCINLSKGA | AKLKSFLLFL**F**---------A |
| E | Effrenium voratum | F | CAJ1456987 | ---------------EAAKAS- | -PRKSP--KA**E**AQDVE--SKD | -SQTP-N-------EIEEPLK | IDEKAAFDLV**L**GNCINLSKGA | ERQIDVVLY--GAQMALAWRL |
| A | Symbiodinium pilosum | F | gene5801 | --------E-**PK**---KEEEP-- | -PQEVP--QA**G**---------- | VCQTP-----**E**EEEPTEEPLR | IDDKAAGDLM**L**GNCINLSKGA | ERQIDIVLYE**C**---------A |
| A | Symbiodinium natans | F | gene5799 | --------S-**PK**---KEAKA-- | -PQEIP--KD**D**---------- | VVQTPL---D**E**DLEDLEVPLR | IDDKAAGDLV**L**GNCVNLSKGA | ERQIDIVLYE**C**---------G |
| A | Symbiodinium tridacnidorum | S | gene20042 | TEN--SDGS-**EV**---KEAETSQ | -PQEVPQHQQ**T**PQET---AQE | ITQTP-----**L**EDMETEEPLR | IDDKAAGDLV**R**GNCINLSKGA | ERQIDIVLYE**H**---------A |
| A | Symbiodinium linucheae | S | gene10151 | PEN--SGGSA**EA**KEKKEAETSE | PPPETP--QQ**T**P------TQE | ITQTP-----**L**EDTETEEPLR | IDDKAAGDLV**R**GNCINLSKGA | ERQIDIVLYE**H**---------A |
| A | Symbiodinium necroappetens | S | gene13677 | AEKSGSGGSA**EA**---KEAETSQ | -PQEAP--QE**T**PQETQE---- | ITQTP-----**L**ED-ETEEPLR | IDDKAAGDLV**R**GNCINLSKGA | ERQIDIVLYE**H**---------A |
| A | Symbiodinium microadriaticum 04 | S | gene21048 | AEKSGSGGSA**EA**---KEAEASQ | -PQETP--QE**T**PQETQEETQG | ITQTP-----**L**ED-ETEEPLR | --------------------- | --------------------- |
| A | Symbiodinium microadriaticum | S | gene36738 | AEKSGSGGSA**EA**---KEAEASQ | -PQETP--QE**T**PQETQEETQG | ITQTP-----**L**ED-ETEEPLR | IDDKAAGDLV**R**GNCINLSKGA | ERQIDIVLYE**H**---------A |
