## Supplementary figures and images for "Positive selection of a starch synthesis gene and phenotypic differentiation of starch accumulation in symbiotic and free-living coral symbiont dinoflagellate species"

### Supplementary figure 2

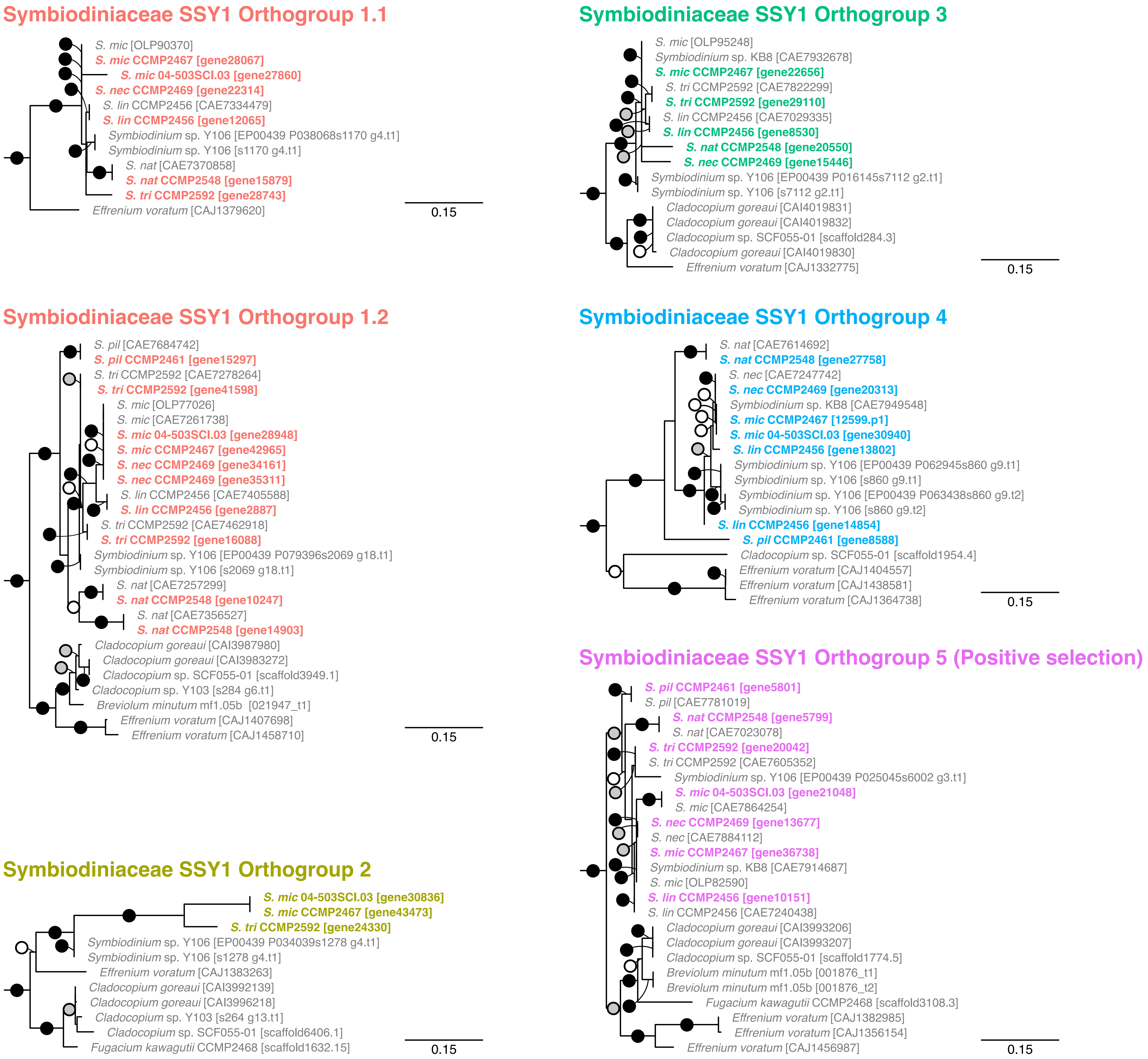

### Supplementary figure 3

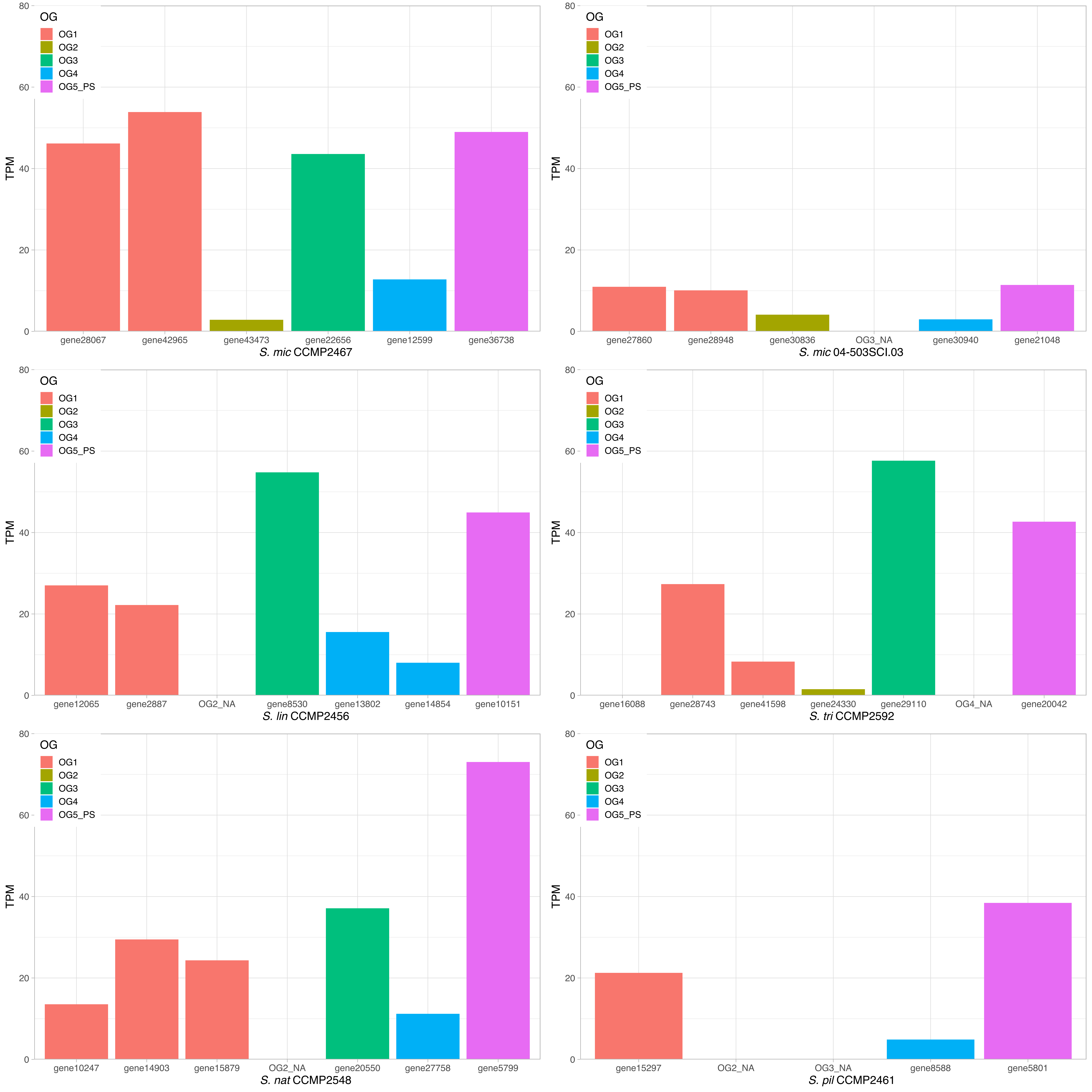

### Supplementary figure 4

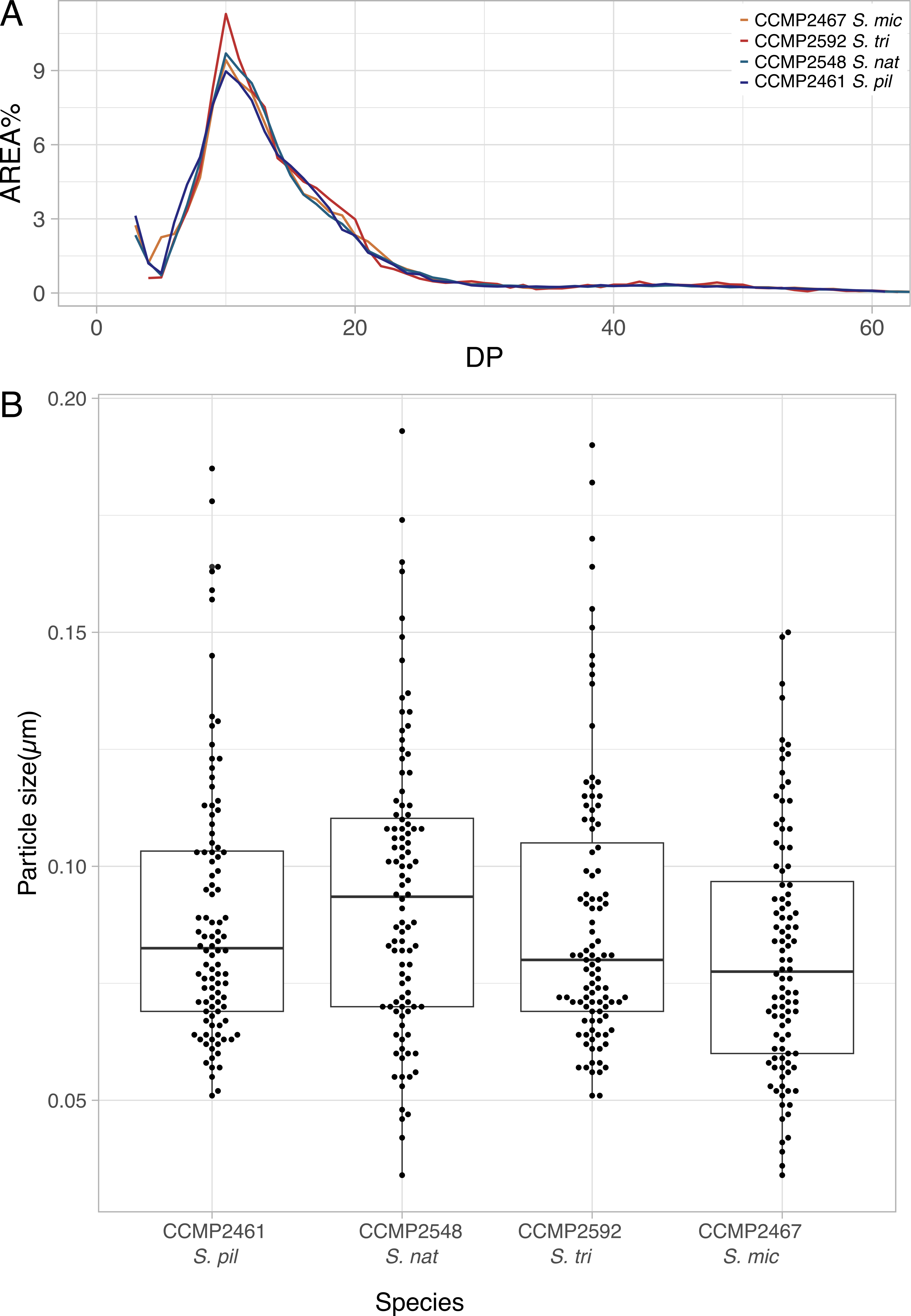
